# Therapeutic potential of SOX9 dysruption in Combined Hepatocellular Carcinoma-Cholangiocarcinoma

**DOI:** 10.1101/2024.05.22.595319

**Authors:** Yoojeong Park, Shikai Hu, Minwook Kim, Michael Oertel, Aatur Singhi, Satdarshan P. Monga, Silvia Liu, Sungjin Ko

## Abstract

Combined hepatocellular carcinoma-cholangiocarcinoma (cHCC-CCA) represents a challenging subtype of primary liver cancer with limited treatment options and a poor prognosis. Recently, we and others have highlighted the context-dependent roles of the biliary-specific transcription factor SOX9 in the pathogenesis of liver cancers using various *Cre* applications in *Sox9*^*(flox/flox)*^ strains, to achieve elimination for exon 2 and 3 of the *Sox9* gene locus as a preventive manner. Here, we reveal the contrasting responses of developmental *Sox9* elimination using *Alb-Cre;Sox9*^*(flox/flox)*^ (*Sox9* LKO) versus *CRISPR/Cas9*-based tumor specific acute *Sox9* CKO in SB-HDTVI-based *Akt-YAP1* and *Akt-NRAS* cHCC-CCA formation. *Sox9* LKO specifically abrogates the *Akt-YAP1* CCA region while robustly stimulating the proliferation of remaining poorly differentiated HCC pertaining liver progenitor cell characteristics, whereas *Sox9* CKO potently prevents *Akt-YAP1* and *Akt-NRAS* cHCC-CCA development irrespective of fate of tumor cells compared to respective controls. Additionally, we find that *Akt-NRAS*, but not *Akt-YAP1*, tumor formation is partially dependent on the *Sox9-Dnmt1* cascade. Pathologically, SOX9 is indispensable for *Akt-YAP1*-mediated HC-to-BEC/CCA reprogramming but required for the maintenance of CCA nodules. Lastly, therapeutic elimination of *Sox9* using the *OPN-CreERT2* strain combined with an inducible *CRISPR/Cas9*-based *Sox9* iKO significantly reduces *Akt-YAP1* cHCC-CCA tumor burden, similar to *Sox9* CKO. Thus, we contrast the outcomes of acute *Sox9* deletion with developmental *Sox9* knockout models, emphasizing the importance of considering adaptation mechanisms in therapeutic strategies. This necessitates the careful consideration of genetic liver cancer studies using developmental Cre and somatic mutant lines, particularly for genes involved in hepatic commitment during development. Our findings suggest that SOX9 elimination may hold promise as a therapeutic approach for cHCC-CCA and underscore the need for further investigation to translate these preclinical insights into clinical applications.

## INTRODUCTION

Combined hepatocellular carcinoma-cholangiocarcinoma (cHCC-CCA) represents a rare and intriguing entity in the spectrum of primary liver cancers, characterized by the coexistence of hepatocellular and cholangiocellular differentiation within the same tumor^1-3^. This dual histological phenotype poses significant diagnostic, prognostic, and therapeutic challenges, reflecting the complex interplay between hepatocytic and biliary lineages in liver tumorigenesis^4^. Especially noteworthy is the distinct response of this tumor type to broad-spectrum therapeutic and immune therapies, making therapeutic strategy largely reliant on basic histological observations, such as measuring the ratio of respective tumor types^5^. Despite its clinical and pathological significance, the underlying molecular mechanisms driving the development and progression of cHCC-CCA remain poorly understood.

Traditionally, HCC and CCA are thought to originate from hepatocytes (HCs) and cholangiocytes (biliary epithelial cells; BECs), respectively, as evidenced by typical cellular morphology, unique structures, and expression of cell type-specific markers. However, the frequent detection of human CCA and cHCC-CCA in the pericentral area of the liver lobule, a region anatomically different from native biliary structures, as well as the documented occurrence of HC-to-BEC differentiation in various chronic cholestasis models, has led to speculation that hepatocytes may also be the origin of a subset of human CCA and cHCC-CCA^4^.

Indeed, numerous studies have provided evidence supporting this theory by inducing HC-derived cHCC-CCA using HC-specific co-expression of proto-oncogenes and biliary lineage commitment genes, such as myristoylated *Akt* (*Akt*) and constitutive-active YAP1 or NRAS^6, 7^. Overexpression of these oncogenes successfully produces separate regions of HNF4α^+^;panCK^+^;CK19^-^ poorly differentiated HCC and HNF4α^-^;CK19^+^ CCA, resembling the clinical cHCC-CCA tumor pathology^6-8^.

Recently, Liu et al., reported that the biliary-specific transcription factor SOX9 determines the fate of YAP1 alone-dependent murine liver cancer; chronic deletion of SOX9 in HC suppresses YAP1-mediated CCA-like tumor formation while promotes aggressive HCC tumor development, suggesting SOX9 as a major commitment of YAP1-mediated liver cancer lineage^9^. However, whether SOX9 is required to maintain the biliary fate of CCA within fully developed advanced cHCC-CCA remains elusive. Additionally, whether SOX9 removal converted CCA lesion into HCC fate or simply eliminated CCA tumor during tumor formation thereby remaining HCC only particularly in YAP1 independent cHCC-CCA settings is unknown. Furthermore, given the validated difference in chronic versus acute gene deletion in diseased liver^10^, the discrepancy between chronic and acute SOX9 elimination on HC-driven cHCC-CCA development has not been thoroughly examined.

Herein, we reveal that unlike chronic developmental *Sox9* deletion, acute and therapeutic elimination of *Sox9* reduces overall cHCC-CCA tumor burden in both *Akt-NRAS* and *Akt-YAP1* models while DNMT1 is partially involved in SOX9-dependent maintenance of *Akt-NRAS* cHCC-CCA. These findings underscore the disparate roles of SOX9 in stage-dependent distinct roles in liver cancer, with potential biological and therapeutic implications.

## RESULTS

### Forced expression of AKT and YAP1 in HC yields cHCC-CCA while chronic elimination of *Sox9* induces molecular phenotype switch to an aggressive HCC

Previously, it has been reported that sleeping beauty transposon/transposase-hydrodynamic tail vein injection (SB-HDTVI) delivery of myristoylated *Akt (Akt)* and *YAP1 S127A (YAP1)*, induced HC-derived cHCC-CCA in a Notch-dependent manner^7, 8^. Tumor-specific *Notch2* deletion switched the tumor type in the *Akt-YAP1* model from cHCC-CCA to a benign hepatocellular adenoma-like tumor at the expense of CCA^7^. Moreover, HC-specific *Sox9* removal prevent YAP1 alone-mediated HC transformation into BEC-like HCC while provokes pure but aggressive HCC indicating *Sox9* is critical for YAP1-driven HC plasticity into biliary lineage^9^. Given that *Sox9* is a well-established direct target of NOTCH2^11^, to investigate the roles for *Sox9* in the lineage commitment of cHCC-CCA, we first sought to examine the impact of cell autonomous *Sox9* deletion, in the *Akt-YAP1*-driven cHCC-CCA tumorigenesis. *Akt* and *YAP1* plasmids were co-delivered by SB-HDTVI into the *Alb-Cre;Sox9*^*flox/flox*^ (*Sox9* LKO) or *Sox9*^*flox/flox*^ (LWT) in which *Sox9* is deleted HC and BEC initiated around E15 days^12^, indicating chronic deletion prior to tumor formation (Fig.1A). Notably, *Akt-YAP1* in *Sox9* LKO led to significantly decreased survival, a much greater and lethal tumor burden as seen by significantly greater LW/BW and macroscopically, as compared to the *Akt-YAP1* in LWT mice (Fig.1B-D). Microscopically, in a representative tiled image of a lobe, the *Sox9* LWT livers in *Akt-YAP1* model showed many intensely CK19-positive CCA nodules scattered throughout (Fig.1E). At higher magnification, the tumors were mixed and showed CCA components which was positive for SOX9, YAP1 and CK19 and HCC component with nuclear HNF4α (Fig.2B). However, the entire *Akt-YAP1 Sox9* LKO livers were full of circumscribed tumor foci which were negative or very week for CK19 (Fig.1E). At higher magnification, the tumors were poorly differentiated HCCs, which lacked SOX9, expressed HNF4α and showed scattered and small subset of YAP1 and panCK-positive cells, displaying liver progenitor cell (LPC) characteristics (Fig. 2B). Quantification of CK19 staining in the tiled image (Fig.1E) verified significant decrease in CK19-positive area in *Sox9* LKO as compared to the LWT (Fig.1G). Further, tumors in *Sox9* LKO *Akt-YAP1* model revealed significantly lower gene expression of biliary markers including *Krt19* and *Epcam* in addition to expected loss of *Sox9*, and simultaneous and significant (or almost significant) increases in HC-specific *Hnf4α* and *Tryptophan 2,3-Dioxygenase (Tdo2)* expression when compared to LWT, also suggesting notable HCC at the expense of CCA (Fig. 2C&D). Together, these data support the role of *Sox9* in commitment to CCA-phenotype in the *Akt-YAP1*-driven cHCC-CCA tumorigenesis. Next, we examined cHCC-CCA (n=32) in the available patient TMA, for SOX9 and YAP1 localization. While majority of the mixed tumors were concurrently positive for both markers (28/32 or 87.5%), a small subset was positive for YAP1 but negative or low for SOX9 (4/32 or 12.5%) (Fig.2E). While the numbers are too low to determine impact on overall tumor behavior or prognosis in these cases, this observation does lend clinical credibility to our preclinical observation.

**Figure 1.**
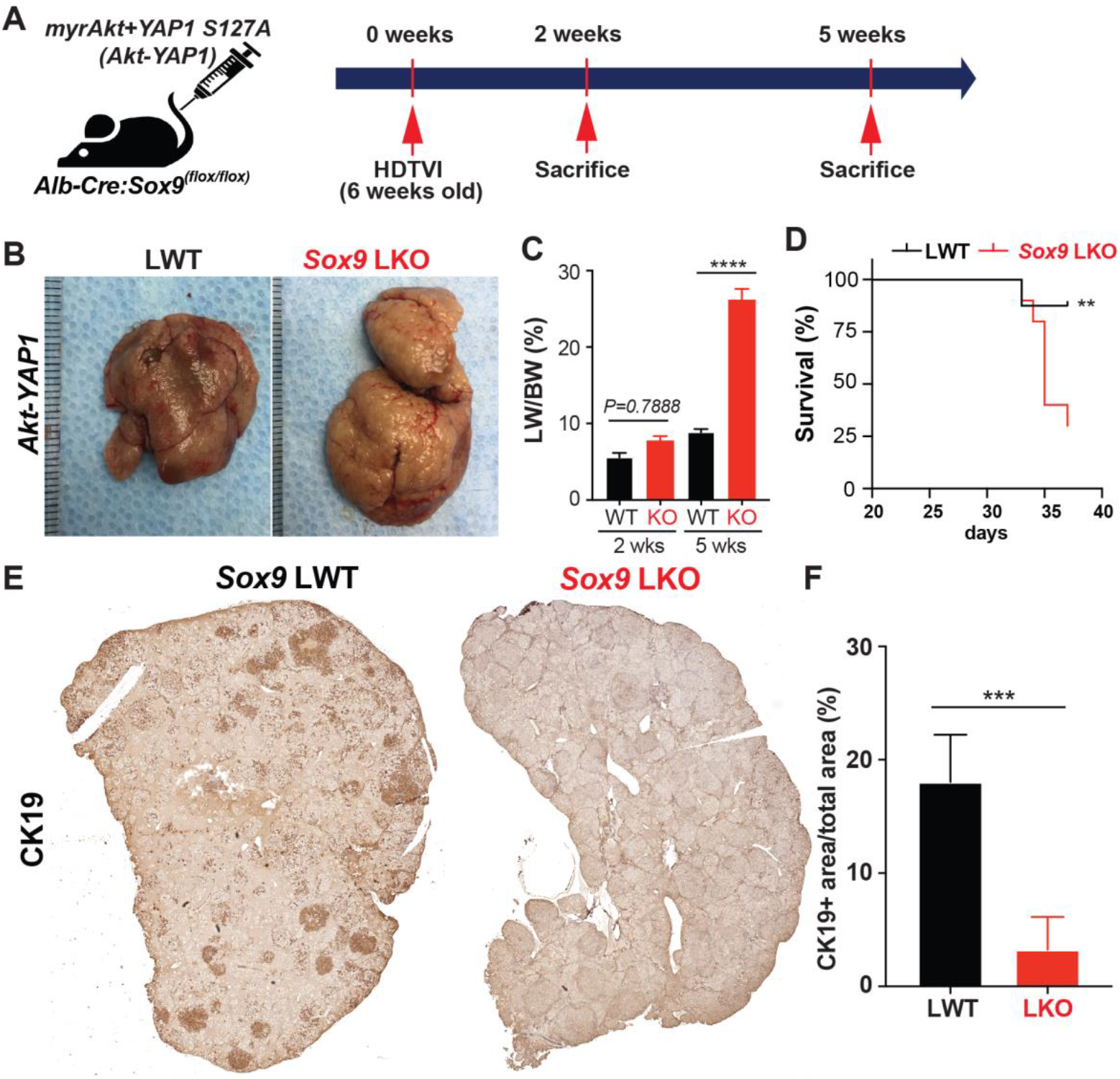
Chronic developmental deletion of *Sox9* switches the fate of *Akt-YAP1*-driven cHCC-CCA to aggressive HCC at the expense of CCA. **(A)** Experimental design illustrating plasmids used for HDTVI, mice used in study and time-points analyzed. **(B)** Representative gross images from *Akt-YAP1*-injected *Sox9*-floxed mice (LWT) and *Akt-YAP1*-injected *Alb-Cre;Sox9*^*(f/f)*^ liver-specific *Sox9* knockout or *Sox9 LKO* mice showing tumor-laden enlarged livers in both cases. **(C)** LW/BW ratio depicts significantly lower tumor burden in *Akt-YAP1 Sox9* LKO as compared to LWT at 5 weeks but not earlier 2 week time point. **(D)** Kaplan–Meier survival curve showing significant decrease in survival of *Akt-YAP1* mice that were *Sox9 LKO* as compared to LWT. **(E)** Representative tiled image of tumor-bearing livers at 5 weeks in *Akt-YAP1* LWT stained for CK19 IHC showing CCA component of the cHCC-CCA staining positive. *Sox9 LKO* livers were full of circumscribed tumors that were negative for CK19 at the same time point. **(F)** Quantification of CK19 IHC verifies significantly reduced staining in *Akt-YAP1 Sox9* LKO as compared to *Akt-YAP1* LWT at 5 weeks as shown in E. Error bar: standard error of the mean; **p<0.01; ***p<0.05; ****p<0.0001.

**Figure 2.**
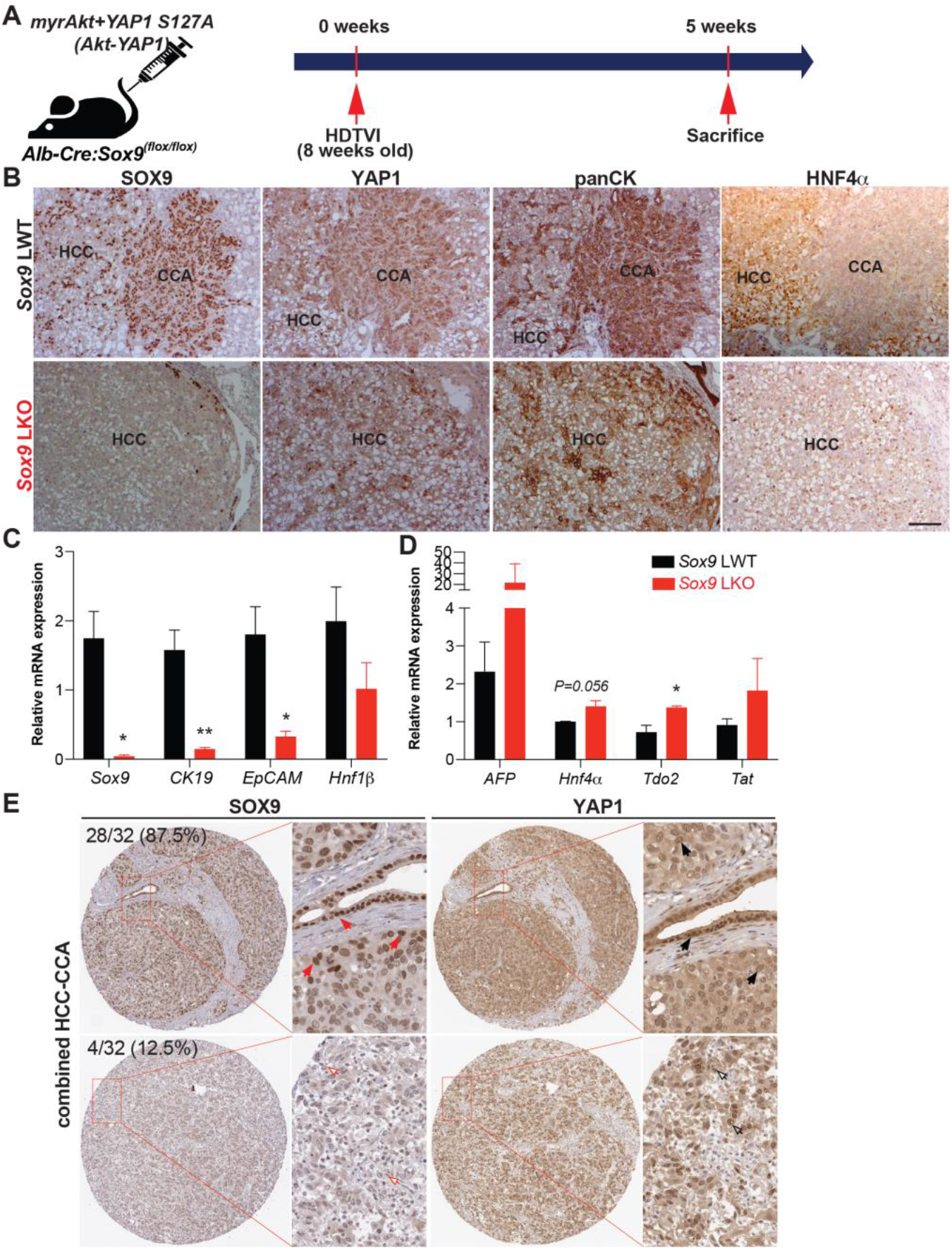
Absence of *Sox9* induces *Akt-YAP1-*mediated HC-driven panCK and HNF4α-positive HCC associated with liver progenitor cell characteristics. **(A)** Experimental design illustrating plasmids used for HDTVI, mice used in study and time-points analyzed. **(B)** Representative serial sections of IHC images of 5-week *Akt-YAP1* LWT show CCA component to be positive for SOX9, YAP1 and panCK and HCC to be strongly positive for HNF4α while in *Akt-YAP1 Sox9* LKO no CCA was seen and HCC was negative for SOX9 but positive for HNF4α with panCK and YAP1 positive cells interspersed in the tumor parenchyma. **(C, D)**. qPCR data showing significantly decreased mRNA expression of *Sox9, CK19 and EpCAM*, and significantly increased expression of *Tdo2* when comparing tumor-bearing livers in *Akt-YAP1* LWT and *Sox9* LKO models at 5 weeks. **(E)**. Representative IHC staining for SOX9 and YAP1 depicting SOX9-low and nucelar YAP1-high or SOX9-high and YAP1-high cHCC-CCA from TMA (32 patients). TMA sections were enlarged for better view on nuclear expression of SOX9 and YAP1. Red arrows point to nuclear SOX9-high cells; black arrows point to nuclear YAP1-high cells; red empty arrows point to nuclear SOX9-negative cells and black empty arrows point to nuclear YAP1-high cells. Percentage of patients positive for each combination are indicated. Scale bars: 100μm; Error bar: standard error of the mean; *p<0.05; **p<0.01.

### Transcriptomic analysis of HCC in *Akt-YAP1* model in *Sox9*-LKO reveals significant similarity to a subset of human HCCs

To directly address the clinical relevance of the HCCs observed in the *Akt-YAP1* model in the absence of *Sox9*, we performed RNA-Seq analysis (GEO accession ID: GSE200472). When comparing the LWT and *Akt-YAP1 Sox9-LKO* livers, 525 genes were upregulated and 199 genes were down-regulated, by FDR=5% and absolute log2 fold change of 1 (Fig.3A). To interpret biological functions of these 724 DEGs, pathway enrichment analysis was performed by Ingenuity Pathway Analysis. Fifty-three pathways were significantly enriched in the *Akt-YAP1 Sox9* LKO livers. To determine if the mouse model mimics a subtype of HCCs in patients, LIHC cohort of TCGA database was analyzed using similar pipeline^13^. When comparing 50 normal or normal adjacent control livers and 374 HCCs the DEGs were enriched in 59 pathways. Ten pathways were commonly altered in mouse and human tumors (Fig.3B). To directly compare mouse and human study, the 724 DEGs from the *Akt-YAP1 Sox9* LKO livers were converted to human homologous genes by the Mouse Genome Database^14^. Human expression data had 546 of the 724 DEGs, and using abs(log2FC)>1 and FDR<0.05, 118 of the 546 DEGs were referred to as the *Akt-YAP1 Sox9* LKO (null) or AYSn signature and applied to the TCGA HCC (Fig.3C). These genes could clearly separate the human normal (orange bar) and HCC (light green bar) groups very well. Lastly, NTP analysis was performed using the *Akt-YAP1 Sox9* LKO signature^15^. In the TCGA cohort, the AYSn signature captured 12% of HCC. This subgroup of patients is enriched in S1 class^16^, (28/46 vs 82/328 in rest of patients, p=0.0015) and an CCA-like signature^17^, (22/46 vs 102/328, p=0.03) (Fig.3D). Altogether, the HCC in *Akt-YAP1 Sox9* LKO model recapitulates a subset of human HCC.

**Figure 3.**
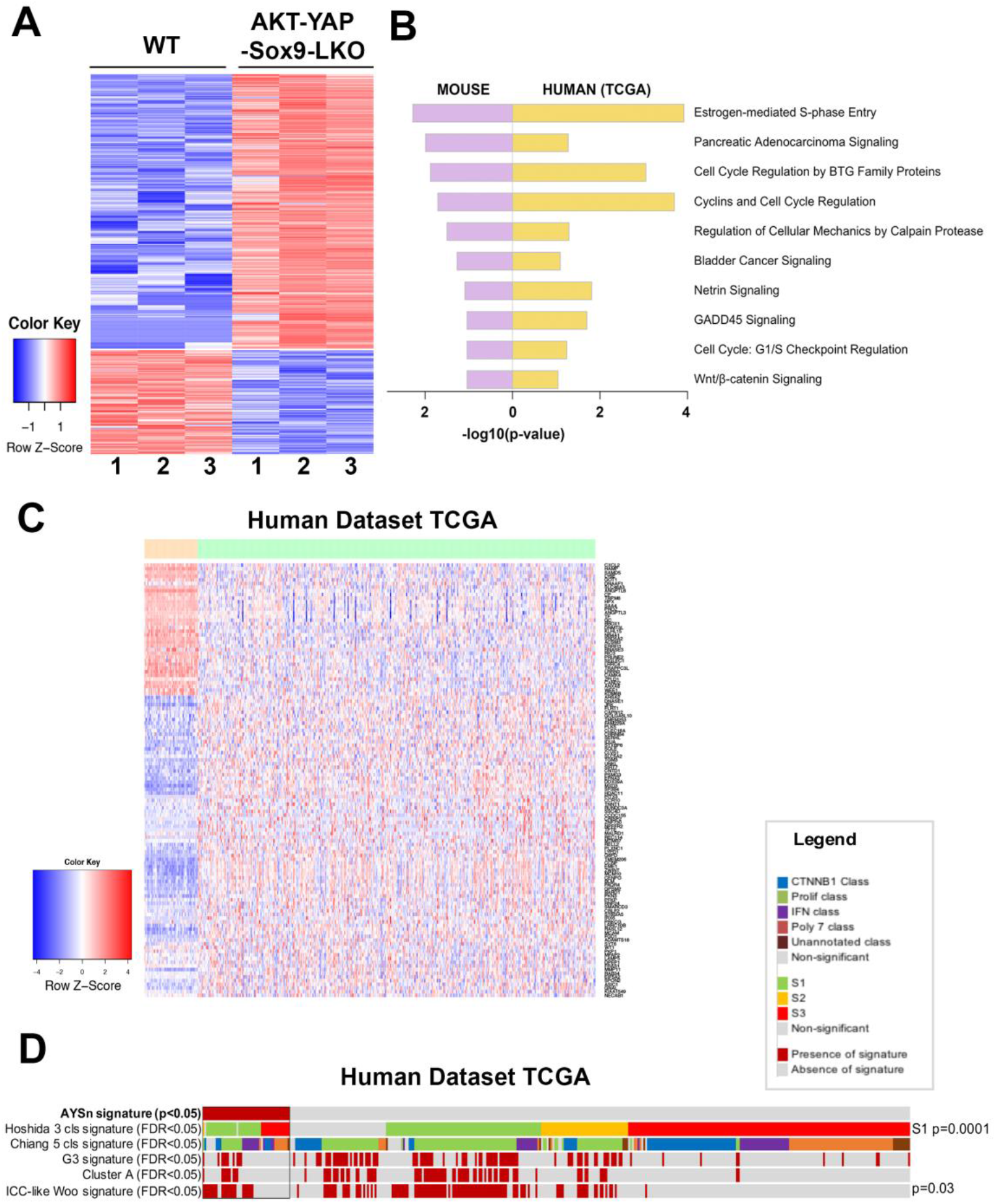
RNA-seq analysis of mouse models and comparison with human liver cancer studies. **(A)** Heatmap for the differentially expressed genes comparing LWT and *Akt-YAP1 Sox9* LKO model. **(B)** Common top enriched pathways between mouse (*Akt-YAP1 Sox9* LKO) and human (TCGA, LIHC) study. **(C)** Heatmap of gene signatures in the TCGA LIHC that are selected by the mouse model (LWT VS *Akt-YAP1 Sox9* LKO). **(D)** Nearest Template Prediction (NTP) analysis of the TCGA LIHC whole-tumor gene expression dataset using the *Akt-YAP1 Sox9* LKO signature (AYSn signature) captured 12% of HCC. This subgroup of patients is enriched in S1 class Hoshida et al, Cancer Research, 2009 (p=0.0015), and an ICC-like signature Woo et al, Cancer Research, 2010 (p=0.03).

### SOX9 is dispensable for *Akt-YAP1-*mediated CCA-like tumor development but required for its maintenance

To investigate whether SOX9 is required for *Akt-YAP1*-driven HC-to-CCA lineage reprogramming, we carefully examined the liver histology in both LWT and *Akt-YAP1 Sox9* LKO mice using serial section IHC to detect HC and BEC markers, including HA-tag (AKT), YAP1, SOX9, HNF4α, and panCK, at the 2 week post HDTVI when fate transition is observed prior to clonal expansion of transduced cells (Fig.4)^8^. Importantly, CCA-like nodules with *Akt-YAP1* transduction (HA-tag^+^; nuclear YAP1^+^) and the retention of intermediate LPC morphology, along with co-expression of HC marker HNF4α and BEC marker panCK, were successfully developed in both LWT and *Sox9*-LKO livers whereas the presence or absence of SOX9 in these nodules differed between the respective livers (Fig.4 black dash lined). This data suggests that while SOX9 is not necessary for YAP1-driven biliary reprogramming, it is essential for the survival and maintenance of these CCA-like tumor nodules. This observation suggests that SOX9 may have stage-dependent distinct roles in fully developed *Akt-YAP1*-CCA at advanced stage, particularly from a therapeutic perspective.

**Figure 4.**
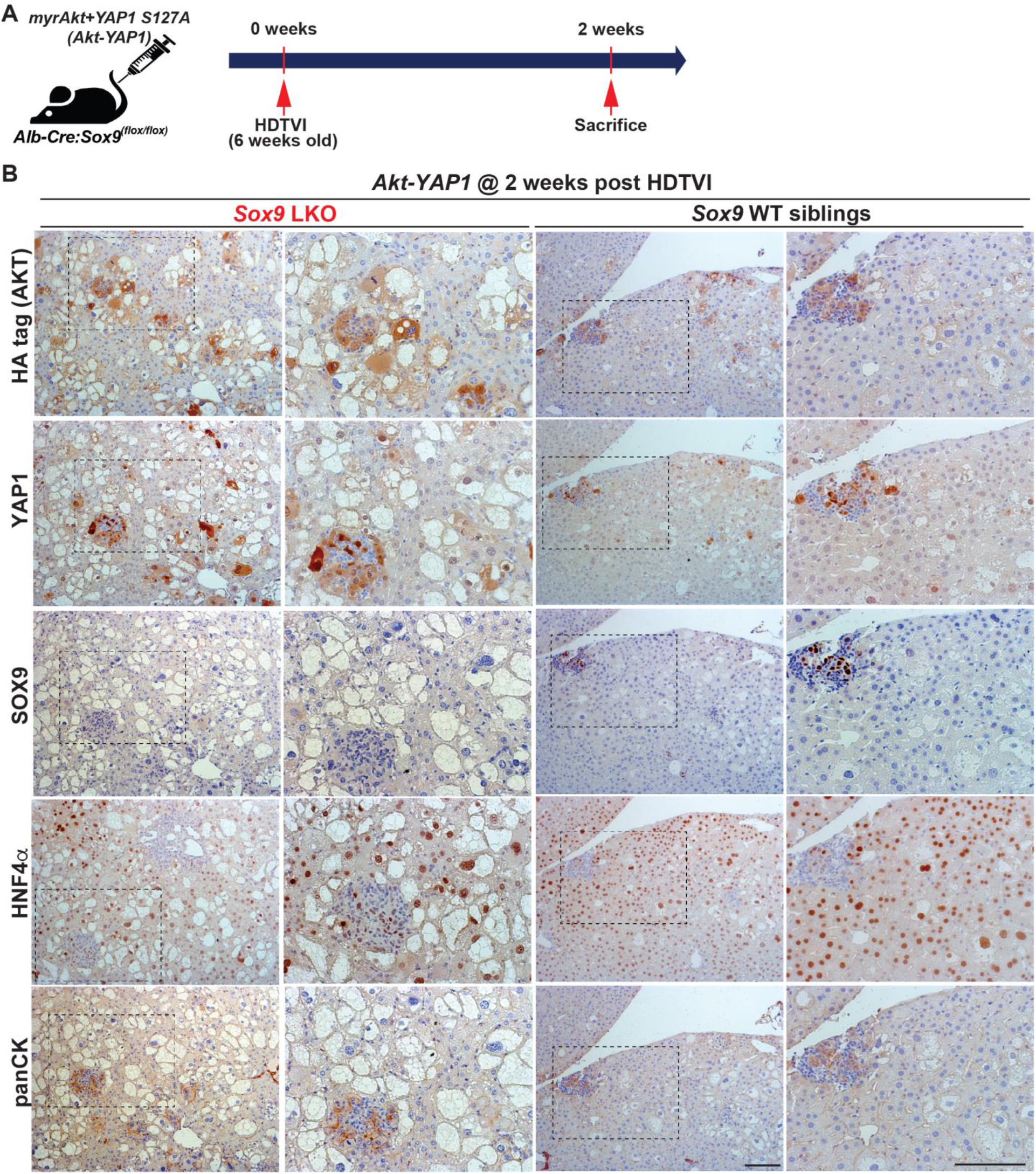
SOX9 is dispensable for *Akt-YAP1*-mediated HC reprogramming into CCA. **(A)** Experimental design illustrating plasmids used for HDTVI, mice used in study and time-points analyzed. **(B)** Representative serial sections IHC images of both *Akt-YAP1* LWT and *Akt-YAP1 Sox9* LKO show CCA-like components to be positive for SOX9, YAP1, panCK and weak positive HNF4α with liver progenitor cell morphology (black dash lined) and HCC to be strongly positive for HNF4α at 2 week post HDTVI. Scale bars: 100μm.

### Distinct roles SOX9 in *Akt-YAP1*-driven HCC tumor in regulating proliferation

Since Sox9 deletion decreases survival in *Akt-YAP1* mice with a larger tumor burden, we next sought to evaluate tumor cell death and proliferation through histologic observation. To investigate, we analyzed WT and *Akt-YAP1 Sox9* LKO liver tissue 5 weeks post-HDTVI using immunofluorescence for Ki-67 to assess proliferation and IHC for TUNEL to measure cell viability (Fig.5A). Interestingly, HCC tumors lacking SOX9 showed a significant increase in the number of Ki-67^+^;HA-tag^+^ tumor cells compared to WT (Fig.5B-C) whereas, there was no significant difference in cell death between the absence of SOX9 and WT (Fig.5D-E). These findings suggest that *Sox9* deletion exacerbates *Akt-YAP1*-mediated CCA-like tumors while promoting the proliferation of HCC, leading to a significantly larger tumor burden that impacts survival, implying the distinct role for SOX9 in liver cancer lineage.

**Figure 5.**
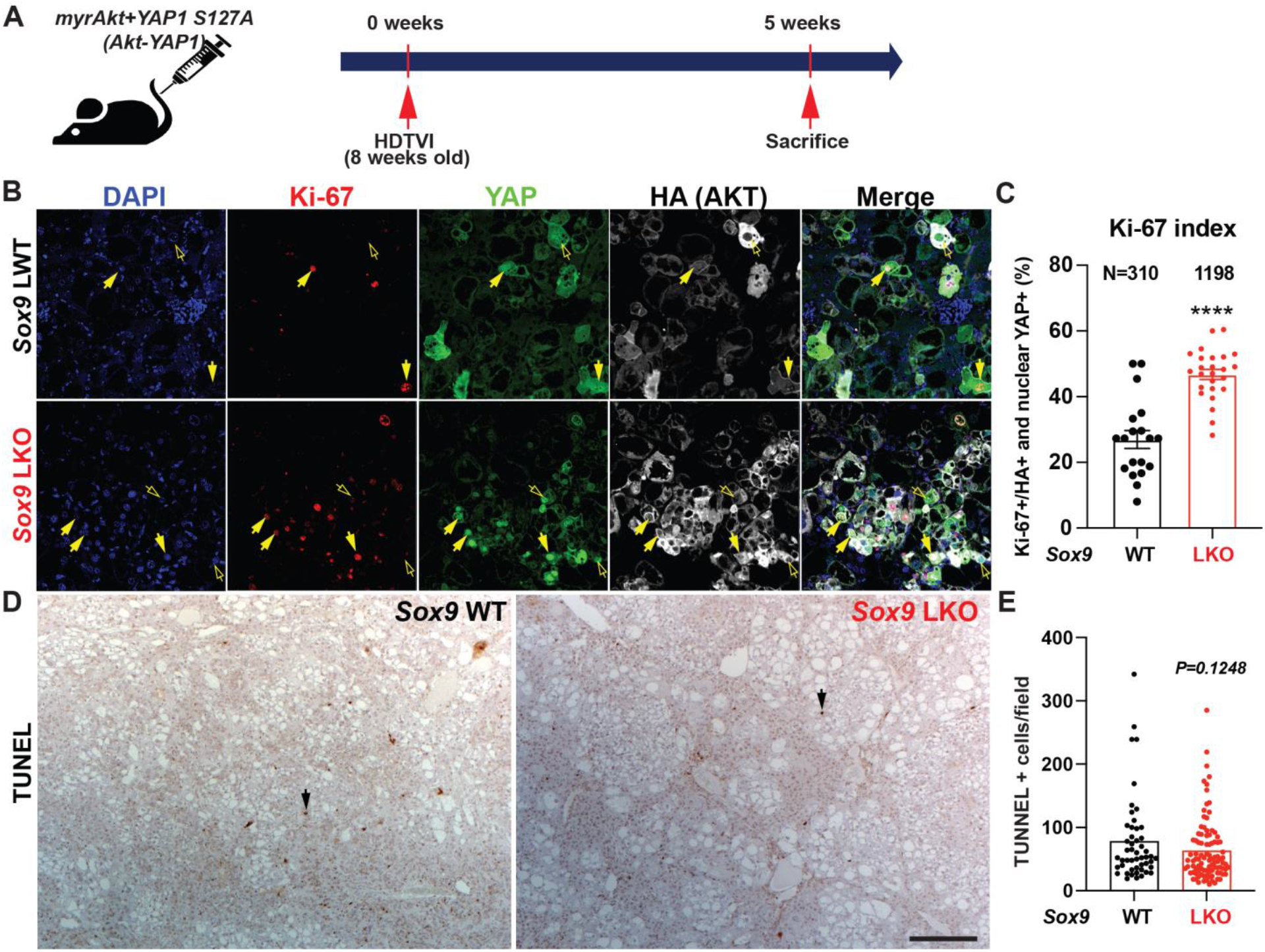
Developmental removal of *Sox9* promotes proliferation of *Akt-YAP1*-mediated liver cancer. **(A)** Experimental design illustrating plasmids used for HDTVI, mice used in study and time-points analyzed. (**B**) Representative IF for Ki-67 (red), YAP1 (green), HA-tag (gray) and DAPI (blue) in liver sections from 5 weeks *Akt-YAP1* LWT or *Sox9* LKO. **(C)** The percentage of Ki-67-positive nuclei normalized to HA-tag-positive total tumor cell nuclei in representative images shown in B, demonstrate significant increase in proliferation of transduced tumor cells in *Sox9* LKO as compared to LWT. **(D)** IHC for TUNEL to detect non-viable tumor cells, shows comparable cell death was evident between *Sox9* LKO and LWT in *Akt-YAP1* model at 5 week time-point. Black arrows indicate nuclear TUNEL-positive apoptotic cells. **(E)** The number of TUNEL-positive nuclei normalized to filed in representative images shown in C, demonstrate comparable cell death in *Sox9* LKO as compared to WT. Scale bars:100 μm; error bar: standard error of the mean; ****p<0.0001.

### Tumor-specific acute *Sox9* loss repress YAP1 or NRAS-dependent cHCC-CCA development

Recently, there have been several publications demonstrating phenotypic differences between chronic developmental gene deletion in the liver using the *Albumin (Alb)*-Cre strain and acute gene deletion mediated by *AAV8-Tbg-Cre/CRISPR-Cas9* system, suggesting an adaptation of HC and BEC against developmental gene deletion. Notably, *Sox9* deletion in HC and BEC since E15 days using *Alb-Cre;Sox9*^*(f/f)*^ animals exhibits delayed bile duct formation^18, 19^, indicating compensatory mechanism, suggesting a potential disparity between acute *Sox9* elimination and possessing developmental adaptation in *Alb-Cre;Sox9*^*(f/f)*^ animals^18, 19^. These observations prompted us to investigate the effect of *Sox9* deletion concurrently with oncogenic events in malignant transformation of HCs. To address this, we utilized an SB-based *CRISPR/Cas9*-mediated inducible *Sox9* deletion vector system applicable via HDTVI delivery (Fig. 6B; manuscript submitted). We employed two validated HC-driven cHCC-CCA models driven by two different combinations of oncogenes. We delivered *SB-STOP*^*(flf)*^*-Cas9-sg-Sox9* and *Cre* expression plasmids to specifically eliminate *Sox9* in tumors along with *Akt-YAP1* or *Akt-NRAS* to induce cHCC-CCA in the presence (*Sox9* CWT) or absence of *Sox9* (*Sox9* CKO). Additionally, based on our previous publication demonstrating the indispensable roles for DNMT1 in biliary fate commitment of Notch or YAP1-mediated liver cancer formation, we delivered a plasmid expressing full-length *Dnmt1* into *Sox9* CKO to investigate its association with SOX9 during liver cancer development. We carefully evaluated tumor formation and examined tumor characteristics at 5 weeks post HDTVI (Fig.6A). All CWT mice developed a significant burden of liver cancer, with liver LW/BW reaching 30 for *Akt-NRAS* (Fig.6C) and 8 for *Akt-YAP1* (Fig.6D) animals, respectively. Remarkably, *Cas9*-mediated acute *Sox9* disruption significantly suppressed both *Akt-YAP1* and *Akt-NRAS*-driven cHCC-CCA development regardless of *Dnmt1* delivery, as all *Sox9* CKO mice remained healthy at 5 weeks when they were sacrificed for comparison to CWT mice. Grossly, *Akt-YAP1 Sox9* CKO or *Akt-NRAS Sox9* CKO mice showed only rare tumors and significantly lower LW/BW ratios compared to the widespread gross disease in CWT (Fig.6C-F). Microscopic observation also depicts the significant decrease of HA-tag^+^ *Akt-NRAS* and *Akt-YAP1* cHCC/CCA tumors in *Sox9* CKO liver supporting gross and LW/BW observations (Fig.6G,I). However, *Dnmt1* re-expression in *Sox9* CKO liver slightly but significantly restored tumor formation driven by *Akt-NRAS* (Fig.6H), but not *Akt-YAP1* (Fig.6I), implying the partial involvement of *Dnmt1* in *Akt-NRAS* tumor development under the SOX9. Importantly, SOX9 IHC confirmed effective elimination both in *Sox9* CKO and *Sox9* CKO-*Dnmt1* livers, indicating successful *Sox9* deletion using our plasmid system. Together, in contrast to *Alb-Cre* strain-driven developmental *Sox9* removal, tumor-specific acute *Sox9* elimination robustly prevents *Akt-YAP1* or *Akt-NRAS*-mediated cHCC/CCA formation irrespective of the lineage of liver cancer, while *Dnmt1* is partially responsible for *Sox9* contribution in *Akt-NRAS* liver cancer development. This data may imply the existence of adaptive compensation of HCs against liver -specific developmental deletion of *Sox9*, which induces a distinct response against *Sox9* elimination, necessitating the examination of therapeutic potential of *Sox9* targeting at an advanced stage.

**Figure 6.**
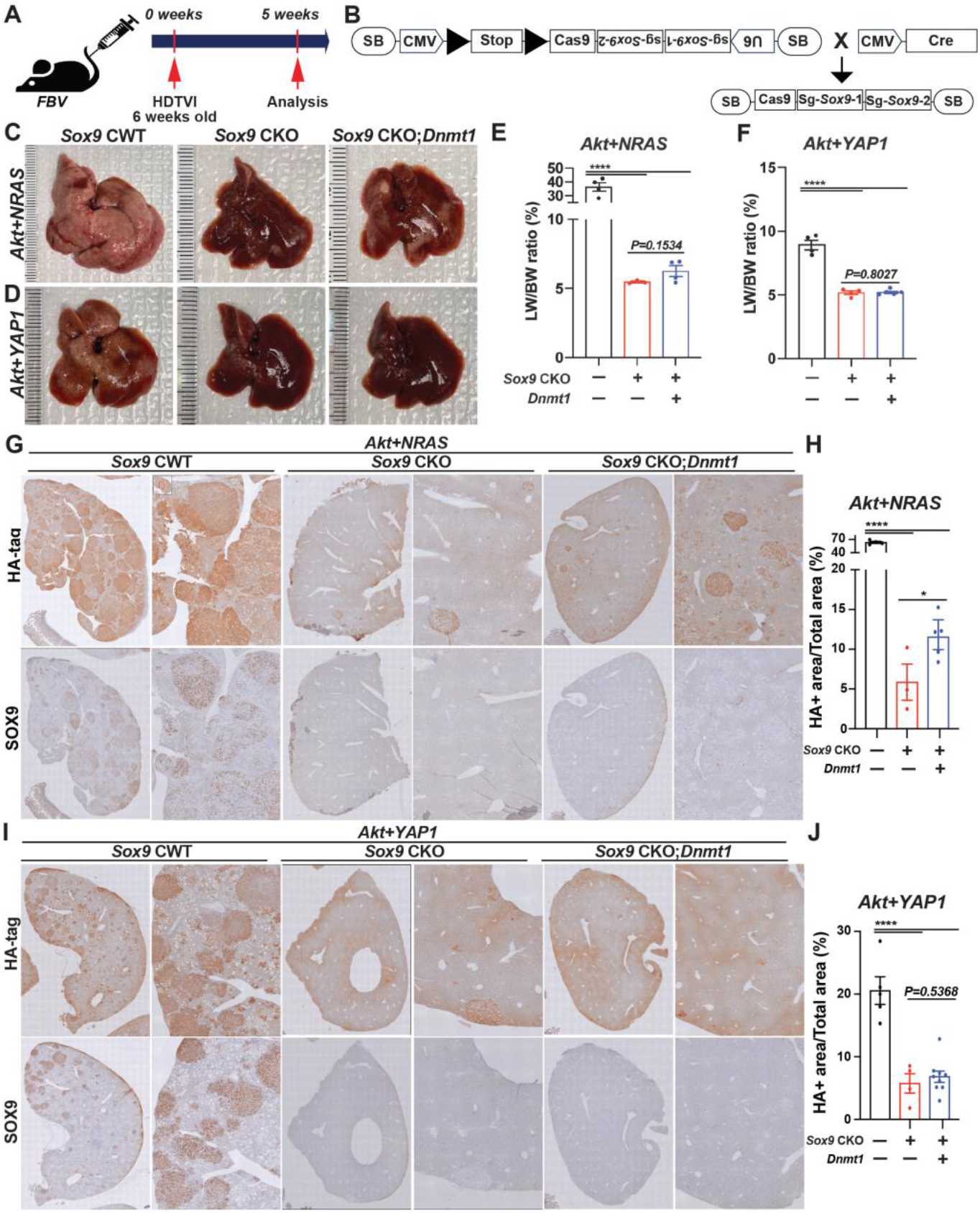
Acute *Sox9* deletion prevent the formation of combined HCC-CCA mediated by *Akt-YAP1* or *Akt-NRAS*, although *Akt-NRAS* tumor exhibit partial dependence on *Dnmt1*. **(A)** Experimental design illustrating plasmids used for HDTVI, mice used in study and time-points analyzed. **(B)** A model illustrating the experimental design utilizing Sleeping Beauty transposon/transposase-CRISPR/Cas9-based inducible *Sox9* knockout plasmid. Representative gross images from *Akt-NRAS* **(C)** or *Akt-YAP1* **(D)**-injected WT (CWT), acute *Sox9*-knockout (CKO) and *Dnmt1*-injected *Sox9-CKO* (*Sox9* CKO-*Dnmt1*) livers showing tumor burden. **(E)** LW/BW ratio depicts significantly lower tumor burden in *Akt-NRAS Sox9* CKO and *Akt-NRAS Sox9* CKO-*Dnmt1* mice as compared to CWT at 5 weeks. **(F)** LW/BW ratio depicts significantly lower tumor burden in *Akt-YAP1 Sox9* CKO and *Akt-YAP1 Sox9* CKO-*Dnmt1* mice as compared to CWT at 5 weeks. **(G**,**H)** Representative IHC images of tumor-bearing livers at 5 weeks in *Akt-NRAS* CWT, *Akt-NRAS Sox9* CKO and *Akt-NRAS Sox9* CKO-*Dnmt1* liver stained for HA-tag and SOX9 showing cHCC-CCA component. HA-tag^+^ *Akt-NRAS* cHCC/CCA tumor burden was robustly abrogated in *Sox9* CKO livers while slightly but significantly restored in *Dnmt1*-injected *Sox9*-CKO livers. **(I**,**J)** Representative IHC images of tumor-bearing livers at 5 weeks in *Akt-YAP1* CWT, *Akt-YAP1 Sox9* CKO and *Akt-YAP1 Sox9* CKO-*Dnmt1* liver stained for HA-tag and SOX9 showing cHCC-CCA component. HA-tag^+^ *Akt-YAP1* cHCC/CCA tumor burden was robustly abrogated in both *Sox9* CKO and *Sox9* CKO-*Dnmt1* livers. Error bar: standard error of the mean; *p<0.05; ****p<0.0001.

### Therapeutic deletion of *Sox9* significantly reduces advanced *Akt-YAP1* liver cancer

To assess the therapeutic effect of *Sox9* deletion in advanced *Akt-YAP1* or *Akt-NRAS* cHCC/CCA, we employed *Osteopontin (OPN)-CreERT2* strains, a well-validated BEC-specific Tamoxifen (TM)-inducible *Cre* expression system^20^. As previously confirmed by other groups, the administration of triple intraperitoneal (i.p.) injections of 100mg/kg of TM effectively induces the *Cre*-mediated deletion of the *Stop* cassette, along with the floxed forms of *Tdtomato reporter*, as evidenced by the absence of RFP in the corresponding CCA and cholangiocyte (masnucript submitted elsewhere). Given the widespread expression of SOX9 in fully-developed *Akt-YAP1* or *Akt-NRAS* CCA regions as well as poorly differentiated HCC regions (Fig.2B), we delivered *SB-STOP*^*(flf)*^*-Cas9-sg-Sox9* along with *Akt-YAP1* or *Akt-NRAS* plasmid into *OPN-CreERT2* mice to induce liver cancer. We then injected 100 mg/kg of TM (i.p.) 3-4 weeks after HDTVI, for a total of 3 injections, to achieve tumor-specific elimination of *Sox9* when liver cancer was fully developed at an advanced stage (*Sox9* iKO). This was followed by assessments at 6-8 weeks post-HDTVI (Fig.7A). As a control, we injected TM into the WT littermates or coron oil into the *OPN-CreERT2* strains (*Sox9* iWT) that were also injected with the same dose of *Akt-YAP1* or *Akt-NRAS* along with *SB-STOP*^*(flf)*^*-Cas9-sg-Sox9* plasmid and sacrificed at the same stage as the experimental group with *Sox9* iKO mice. Notably, the injection of *Sox9* iKO significantly decreased tumor burden at 6-8 weeks post-HDTVI (Fig.7B-E), indicating the potent therapeutic effect of this treatment regimen. Consistently, the *Akt-YAP1* or *Akt-NRAS* iWT mice developed significant liver cancer with LW/BW reaching 15 and 20 respectively, while there was a significantly lower burden of tumor in the iKO mice at 6-8 weeks, comparable LW/BW with mice without tumors with normal rage 4-5 at the same stages, suggesting a strong therapeutic effect (Fig.7C,E). Notably, a subset of CCA-like tumors in iWT liver displayed SOX9^-^ nodules, suggesting evident leakiness of the floxed Stop cassette in our system, however, still significant and robust reduction of HA-tag^+^ liver cancer in iKO livers implies notable therapeutic potential of *Sox9* elimination in advanced *Akt-YAP1* or *Akt-NRAS* cHCC-CCA tumors (Fig.7F,G).

**Figure 7.**
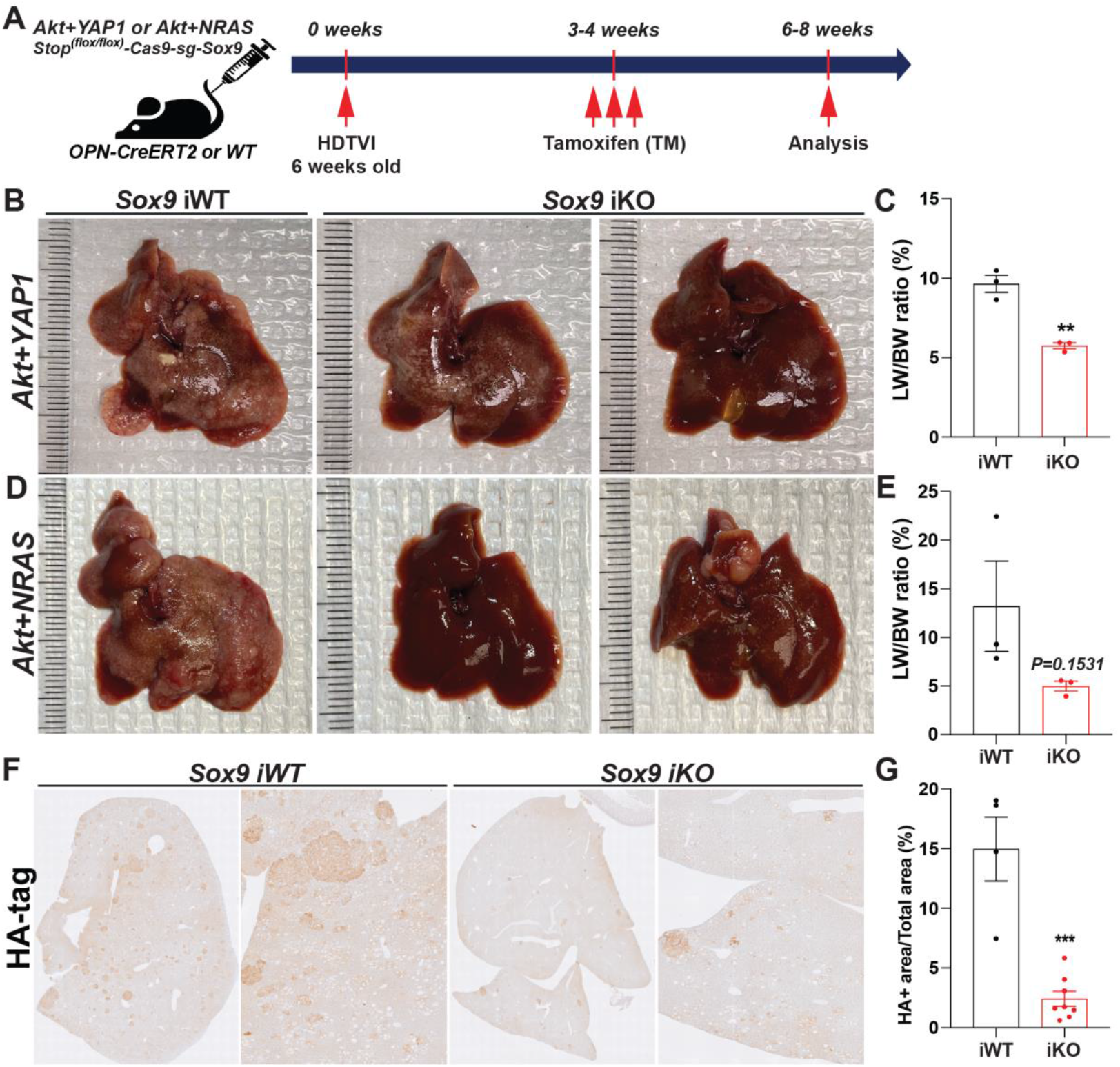
Therapeutic *Sox9* elimination reduces *Akt-dependent* advanced combined HCC-CCA. **(A)** Experimental design illustrating plasmids used for HDTVI, Tamoxifen treatment and mice strain used in study and time-points analyzed. **(B&D)** Representative gross images from *Akt-YAP1* (B) or *Akt-NRAS* (D)-injected *OPN-CreERT2* mice along with *Stop*^*(f/f)*^*-Cas9-sg-empty* (*Sox9*-iWT) and *Stop*^*(f/f)*^*-Cas9-sg-Sox9* (*Sox9*-iKO) treated with Tamoxifen (100 mg/kg triple) displaying gross tumor burden. **(C&E)** LW/BW ratio depicts significantly lower tumor burden in *Akt-YAP1* (C) or *Akt-NRAS* (E) *Sox9* iKO as compared to iWT at 6 weeks **(F)** Representative IHC image of tumor-bearing livers at 6 weeks in *Akt-YAP1* iWT stained for HA-tag IHC showing component of the cHCC-CCA staining. *Sox9 iKO* livers significantly reduced HA-tag^+^ tumor burden. **(G)** Quantification of HA-tag IHC verifies significantly reduced staining in *Akt-YAP1 Sox9* iKO as compared to *Akt-YAP1 Sox9* iWT at 6 weeks as shown in B. Error bar: standard error of the mean; **p<0.01; ***p<0.05.

## DISCUSSION

Primary liver cancer, including HCC and CCA, arises from malignancies of hepatic parenchymal epithelial cells, HC and BEC, derived from the same parental cell, hepatoblast, during development^21^. Interestingly, the liver exhibits a remarkable capacity for regeneration, characterized by the cellular plasticity of these two adult cell populations, involving diverse and complex molecular signaling pathways^21^. Particularly, crucial lineage-specific transcriptional regulatory components such as SOX9, HNF4α, HIPPO-YAP1, and HNF1α/β play important roles in forced cellular reprogramming both *in vitro* and *in vivo*^*21*^. Indeed, several groups have demonstrated the translation of this plasticity into cancer settings, especially in mixed HCC/iCCA and/or cHCC-CCA tumors^8, 9, 22-25^. Mixed HCC-iCCA and/or cHCC/CCA are also evident in clinical practice and the cellular and molecular basis remains unknown^26^. *Akt-YAP1* but not *Akt-NICD* model displayed such combined tumors suggesting distinct roles of YAP1 and Notch signaling in cooperating with AKT activation in hepatobiliary tumorigenesis^8, 24^. Activation of YAP1 appears to be driving a more hepatoblast/LPC-like cell fate which then evolves into either CCA or HCC^24^. Importantly, YAP1 activation in conjunction with active-β-catenin yielded hepatoblastoma in SB-HDTVI model^27^. Additionally, SOX9 was critical in directing cholangiocyte and eventually CCA cell fate in the *Akt-YAP1* model since chronic *Sox9* LKO drove the *Akt-YAP1*-reprogramed cell towards HC at the expense of biliary fate. Indeed, SOX9 has been shown to be essential for proper bile duct differentiation during hepatic development^18, 28^. What was also unexpected was that elimination of *Sox9* in the *Akt-YAP1* models not only prevented development of CCA component of the cHCC-CCA tumors, it led to a more aggressive HCC with higher proliferative index. This suggests that in the context of YAP1 activation, SOX9 may be restricting HC or HCC cell proliferation. This is a novel observation and while *Sox9* has been shown to be both upregulated^29^ or downregulated^30^ by YAP1 signaling and appears context-dependent, how SOX9 restricts YAP1 signaling and suppress cell proliferation in transformed hepatocytes remains unknown. It should be noted that SOX9 upregulation and downregulation have both been observed in various tumors and thus the overall biological outcome may be tissue-dependent^31^. In our study, SOX9 has dual roles of not just regulating biliary differentiation of hepatoblast to yield CCA, but also restrict YAP1-dependent HC proliferation, such that in its absence, the *Akt-YAP1* driven HCC are more proliferative and aggressive.

A recent study led by Dr. Yang’s group demonstrated two dominant roles for SOX9 in *YAP1* alone-mediated liver cancer: indispensable roles in lineage determinant for HC-to-iCCA formation and responsibility for the severity of *YAP1*-HCC using *AAV8-Cre;Sox9(f/f)*-mediated *Sox9* ablation^9^. However, we previously reported that SOX9 is dispensable in NOTCH-driven HC-to-BEC/iCCA reprogramming, whereas it is involved in tumor cell viability and proliferation in *Akt-NICD* HC-driven iCCA models^8^. This indicates context-dependent roles for SOX9 in fate control and cell viability within the mammalian liver cancer. These observations also suggest that anticipating the roles of biliary factors in clinical liver cancer settings is extremely difficult and complex due to the complicated and heterogeneous molecular signature of human liver cancers. In the current study, we mainly used the SB-HDTVI-based *Akt-YAP1* and *Akt-NRAS* HC-driven cHCC-CCA models to investigate the roles of *Sox9* in liver cancer setting including cHCC-CCA. Importantly, some of our observations are similar to those of *YAP1* alone-driven liver cancer study, while a large part of our investigation demonstrates distinct responses, with multiple caveats. Consistent with observations in *YAP1* alone-mediated liver cancer setting, chronic developmental deletion by *Sox9* LKO prior to HDTVI delivery of *Akt-YAP1* induced a switch of tumor lineage from cHCC-CCA to aggressive and poorly differentiated HCC with LPC characteristics, genetically representing a subset of clinical HCC cases. However, in contrast to *Sox9* LKO or YAP1-alone-mediated liver cancer study, acute *Sox9* disruption using the *CRISPR/Cas9* system robustly repressed cHCC-CCA formation driven by two different oncogene drivers, irrespective of tumor fate.

These discrepancies raise several caveats that need to be carefully considered in murine liver cancer models. In particular, the difference between *Sox9 LKO* and *Sox9 CKO* may underscore the importance of hepatic adaptation against deletion of the genes involved in hepatic competence and specification during development, *Sox9* in current case, which may be responsible for the dependency cHCC-CCA formations. From this perspective, comprehensive investigation the adaptive compensatory roles for validated compensation genes will be the pertienet studies noteworthy to elucidate the mechanism behind these differences. Furthermore, the distinct tumor microenvironment between HDTVI delivery of *YAP1* and non-invasive TET-ON *YAP1* expression in HC used for the *YAP1* alone liver cancer model, along with the collaboration of constituvel active AKT, needs to be carefully compared to conclude these different observations.

Previously, we reported that the NOTCH-YAP1-DNMT1 axis plays indispensable and permissive roles in HC reprogramming into CCA whereas DNMT1 is dispensable for the maintenance of CCA^8^. Given that *Sox9* LKO specifically abrogates the *Akt-YAP1* CCA region, similar to pharmacologic DNMT1 inhibition, we posited an association between SOX9 and DNMT1 in CCA fate commitment in our cHCC-CCA models. Interestingly, SOX9-DNMT1 is partially involved only in *Akt-NRAS* tumor formation but not in *Akt-YAP1* tumor development. This may suggest that DNMT1 and SOX9 are independently regulated under YAP1-provoking HC-originated CCA tumor region, while NRAS-SOX9-DNMT1 signaling cascade may be active in *Akt-NRAS* CCA tumor cells. However, further functional validation, including examining the association of YAP1 or SOX9 with pharmacologic/genetic DNMT1 regulation on *Akt-NRAS* CCA, will be an interesting focus for future studies.

Importantly, we observed the successful formation of SOX9^-^ CCA-like nodules in *Akt-YAP1 Sox9* LKO livers at an early stage at 2 weeks post HDTVI, expressing equivalent markers to SOX9^+^ CCA-like nodules in *Sox9* WT livers, which is similar with case of *Akt-NICD Sox9* KO livers forming SOX9^-^ *AKT-NICD* CCA nodules. This indicates the dispensable roles of SOX9 in both *Akt-YAP1* or *Akt-NICD*-driven HC-to-BEC/CCA reprogramming, whereas SOX9 is required for the maintenance of *Akt-YAP1* but not *Akt-NICD* CCA tumors. This supports our claim of overlapping and/or context/stage-dependent roles for SOX9 in liver cancer including CCA while this observation need to be carefully repeated in our *Sox9* CKO models to exclude the effect of hepatic adaptation in *Akt-YAP1* CCA formation.

Lastly, considering the significant therapeutic effect of *Sox9* ablation in advanced cHCC-CCA, similar to our *Sox9* CKO (acute deletion) data but contrasting with *Sox9* LKO (developmental deletion) in a preventive manner, there is a need to revisit tumor studies conducted using developmental *Cre* strains-mediated gene KO systems, which may overlook adaptation. This suggests the necessity for future studies to reevaluate therapeutic potentials, anticipating distinct response impact information as preclinical data translates into relevance for human cancer patients.

## Materials and Methods

### Mouse husbandry and breeding

All animal care and experiments were performed in accordance with the Institutional Animal Care and Use Committee (IACUC) at the University of Pittsburgh. *Sox9*^*(flox/flox)*^ and *Yap*^*(flox/flox)*^ mice were purchased from Jackson Laboratories for breeding. All transgenic and KO mouse lines were maintained on the immunocompetent C57BL/6 genetic background. All animals ranged from 8-12 weeks in age for analysis and were from either gender.

### Patient data

Study approval for all human tissue samples was obtained from the University of Pittsburgh (IRB# STUDY19070068). All samples were provided by the Pittsburgh Liver Research Center’s (PLRC’s) Clinical Biospecimen Repository and Processing Core (CBPRC), supported by P30DK120531.

TMAs were constructed from archival formalin-fixed paraffin-embedded tissue blocks from 108 cholangiocarcinoma patients seen at the University of Pittsburgh Medical Center and were also obtained from PLRC’s CBPRC supported by P30DK120531. All tumor hematoxylin and eosin (H&E) stained slides were reviewed, and representative areas were carefully selected for tissue microarray construction. Two, random 1.0 mm-sized cores were punched from each patient’s tumor and harvested into recipient blocks. The demographics and additional information of these cases are included in Supplementary Table 2. The TMA were stained manually using antibody against SOX9 (EMD Millipore), YAP (Cell Signaling) as described in IHC sections. Whole slide image capture of the tissue microarray was acquired using the Aperio XT slide scanner (Aperio Technologies). The staining was evaluated and scored by anatomic pathologist (A.S.). Staining for SOX9 and YAP was scored either as 0 (negative), 1+ (mostly cytoplasmic staining or very weak staining in CC tumor cells), or 2+ (strong positive nuclear staining in CC tumor cells). For SOX9 and YAP1, the scores for different tissue sections from each patient were averaged to get a single score per patient (Supplementary Table 2). Mean scores greater than or equal to 1.5 were considered “HIGH” and mean scores less than 1.5 were considered “LOW/NEGATIVE”.

## Please see Online Supplement for Additional Methods

### 1. SUPPLEMENTARY METHODS

#### Animal Models of Intrahepatic Cholangiocarcinoma

The constructs used for mouse SB-HDTVI, including *pT3-EF1α, pT3-EF1α-myrAkt-HA* (mouse), *pT3-EF1α-YAP1 S127A* (human), *pT3-EF1α-NRAS V12 caggs, pCMV-empty, pCMV-Cre*, and *pCMV-sleeping beauty transposase* (SB) were generated or have been described elsewhere ^7, 32, 33^. All the plasmids used for *in vivo* experiments were purified using the Endotoxin Free Maxi Prep kit (Sigma-Aldrich). Six-to-eight weeks old mice were randomized into groups and subjected to the sleeping beauty transposon-transposase and hydrodynamic tail vein (SB-HTVI) protocol as described previously ^32^. Briefly, 10 μg *pT3-EF1α-myrAkt-HA, pT3-EF1α-YAP1 S127A* or 10 μg *pT3-EF1α-myrAkt-HA* and 20 μg of *pT3-EF1α-NRAS V12 caggs*, and 40 μg of *pCMV-Cre (or pCMV-empty), and/or newly generated CMV-LSL-Cas9;U6-sg-Sox9* plasmid (described below) along with the transposase in a ratio of 25:1 were diluted in 2 ml of normal saline (0.9% NaCl), filtered through 0.22μm filter (Millipore), and hydrodynamically injected into the lateral tail vein of mice. All animals were sacrificed between 2-6 weeks of plasmids injections unless otherwise indicated.

#### Patient data

All human tissue samples were provided by Pittsburgh Liver Research Center’s (PLRC’s) Clinical Biospecimen Repository and Processing Core supported by P30DK120531 under approved Institutional Review Board STUDY19070068. TMAs were constructed from archival formalin-fixed paraffin-embedded tissue blocks from 108 cholangiocarcinoma patients seen at the University of Pittsburgh Medical Center and were also obtained from PLRC’s CBPRC supported by P30DK120531. All tumor hematoxylin and eosin (H&E) stained slides were reviewed, and representative areas were carefully selected for tissue microarray construction. Two, random 1.0 mm-sized cores were punched from each patient’s tumor and harvested into recipient blocks. The demographics and additional information of these cases are included in Supplementary Table 2. The TMA were stained manually using antibody against SOX9 (EMD Millipore) and YAP1 (Cell Signaling) as described in IHC sections. Whole slide image capture of the tissue microarray was acquired using the Aperio XT slide scanner (Aperio Technologies). The staining was evaluated and scored by anatomic pathologist (A.S.). Staining for SOX9 and YAP was scored either as 0 (negative), 1+ (mostly cytoplasmic staining or very weak staining in CC tumor cells), or 2+ (strong positive nuclear staining in CC tumor cells). For all 2 markers, the scores for different tissue sections from each patient were averaged to get a single score per patient (Supplementary Table 2). Mean scores greater than or equal to 1.5 were considered “HIGH” and mean scores less than 1.5 were considered “LOW/NEGATIVE”. All scores for each individual section are included in Supplementary Table 1.

### Constructs *CRISPR/Cas9-sg-Sox9* transposon vectors and overexpression vectors

We subcloned CMV-LSL-Cas9-P2A-EGFP fragment from the LSL-Cas9-Rosa26TV plasmid (Addgene #61408) by digesting with PacI (NEB# R0547L) and NsiI (NEB# R0127S), followed by purification using a QIAquick PCR & Gel Cleanup Kit (Qiagen# 28506). Next, we amplified a cassette from the CRISPR-SB plasmid (Addgene #177936) with the High Fidelity PCR EcoDry™ Premix (TaKaRa#639280). This cassette included plasmid replication elements, an SB-transposon, and a sgRNA scaffold-linker-U6 promoter. We assembled this cassette with the CMV-LSL-Cas9-P2A-EGFP fragment using the NEBuilder HiFi DNA Assembly Master Mix (NEB#E2621L).

### Cloning CRISPR sgRNA for the target genes

We determined the *CRISPR sg-Sox9* target sequences using the CHOPCHOP online tool (https://chopchop.cbu.uib.no). Following this, we ordered two complementary single-strand oligonucleotides of the target sequence from IDT with specific overhangs: one with a 5’-CACC overhang and the other with a 3’-ACCC overhang. To insert these sgRNA target sequences into construct, we first phosphorylated them using T4 PNK (NEB#M0201S), then denatured and annealed them to form double-stranded oligos. Next, we linearized the CRISPR/Cas9-sgRNA transposon plasmid with BbsI (NEB# R0539L) and inserted the double-stranded oligo using the NEB Quick Ligation™ Kit (NEB#M2200S). To improve our chances of achieving a knock-out, we incorporated an additional CRISPR sgRNA target sequence with sgRNA scaffold-linker-U6 promoter cassette into the final construct. This was done by inserting another double-stranded oligo into the CRISRP-SB plasmid, which had been digested with BbsI (NEB# R0539L). We then amplified this additional sgRNA sequence-linker-U6 promoter cassette and integrated it into the previous *CRISPR/Cas9-sgRNA* transposon plasmid containing the previous CRISPR sgRNA target sequence. This integration utilized the NEBuilder HiFi DNA Assembly Master Mix (NEB#E2621L), with the linearization performed by PacI (NEB# R0547L). For details on all constructs and primers used in this project can be obtained upon request.

#### Immunohistochemistry (IHC)

Mouse liver tissues were fixed for 48 h in 10% neutralized formalin (Fisher Chemicals), transferred into 70% ethanol and then dehydrated and embedded in paraffin. For IHC, formalin-fixed sections were deparaffinized in graded xylene and ethanol and rinsed in PBS. For antigen retrieval, samples were microwaved for 12 min in pH 6.0 sodium citrate buffer (HA-tag, panCK, SOX9) or pH 9.0 Tris-EDTA buffer (p-AKT, CK19), or were pressure cooked for 20 min in pH 9.0 Tris-EDTA buffer (YAP1 and HNF4α). After cooling, samples were placed in 3% H_2_O_2_ (Fisher Chemicals) for 10 min to quench endogenous peroxide activity. After washing with PBS, slides were blocked with Super Block (ScyTek Laboratories) for 10 min. Sections were incubated for overnight at 4C with the primary antibodies (listed in table below). Sections were then incubated with species-specific biotinylated secondary antibodies (EMD Millipore, listed in table below) for 1 h, at room temperature. Sections were incubated with Vectastain ABC Elite kit (Vector Laboratories) and signal was detected with DAB Peroxidase Substrate Kit (Vector Laboratories) followed by quenching in distilled water for 5 min. Slides were counterstained with hematoxylin (ThermoFisher Scientific), dehydrated to xylene (Fisher Chemicals) and coverslips applied with Cytoseal XYL (ThermoFisher Scientific). To assess cell death, TUNEL staining was performed using the ApopTag Peroxidase In situ Detection kit (Milipore, cat# S7100) according to the manufacturer’s instruction.

#### Immunofluorescence

Paraffin embedded liver sections (5 μm thick) were deparaffinized using xylene (Fisher Chemicals) and rehydrated by incubating the slices in ethanol (100% and 95% v/v, each 3×5 min) and washed in PBS. Heat-induced epitope retrieval was performed for 20 min using a pressure cooker with pH 6.0 sodium citrate buffer. Sections were washed in PBS, permeabilized for 5 minutes with PBS/0.3% Triton X and blocked with PBS/0.3% Triton X/10% bovine serum albumin (BSA) for 45 minutes at room temperature. Sections were incubated with primary antibodies in PBS/0.3% Triton X/10% BSA overnight at 4C. At the end of the incubation, sections were washed thrice and incubated with fluorochrome-conjugated secondary antibodies in PBS/0.3% Triton X/10% BSA for 1h at room temperature, then washed in PBS/0.1% Triton X 3 times. Liver sections were mounted using Prolong Gold Antifade w/DAPI (Invitrogen) and pictures were acquired using LSM700 confocal microscope the and Zen Software (Zeiss).

### List of antibodies for IHC and IF in this study

#### Primary Antibodies

Primary Antibodies:
| Target | Species | Dilution | Source | Catalog Number |
| --- | --- | --- | --- | --- |
| HA-tag | Mouse | 1:50 (IHC) | Cell Signaling | CS2367 |
| Pan-CYTOKERATIN (panCK) | Rabbit | 1:200 (IHC) | Dako | Z0622 |
| SOX9 | Rabbit | 1:200 (IF)<br>1:2000 (IHC) | EMD Millipore | Ab5535 |
| YAP1 | Rabbit | 1:100 (IF and IHC) | Cell Signaling | CS14074 |
| HNF4α | Rabbit | 1:100 (IHC) | Cell Signaling | CS3113 |
| Ki-67 | Goat | 1:100 (IF) | Santa Cruz | sc-7846 |
| CK19 | Rat | 1:10 (IF) | DSHB | TROMA III |

#### Secondary Antibodies

**Secondary Antibodies:**
| **Secondary Antibody** | **Species** | **Source** | **Catalog Number** |
| --- | --- | --- | --- |
| Donkey anti-Rabbit IgG Biotin | Donkey | EMD Millipore | AP182B |
| Goat anti-Mouse IgG (H+L) Biotin | Goat | EMD Millipore | AP181B |
| Alexa-Fluor 555 Donkey anti-Rabbit IgG (H+L) | Donkey | Invitrogen | A31572 |
| Alexa-Fluor 488 Donkey anti-Rat IgG (H+L) | Donkey | Invitrogen | A21208 |
| Alexa-Fluor 647 Goat anti-Mouse IgG (H+L) | Goat | Invitrogen | A21236 |

### RT-qPCR analysis

Whole liver was homogenized in TRIzol™ (Thermo Scientific, Cat# 15596026), treated with chloroform, and nucleic acid was precipitated with isopropanol. Cellular DNA was digested with DNA-free™ Kit (ambion, AM1906), and RNA was reverse-transcribed into cDNA using SuperScript® III (Invitrogen, 18080-044). Real-time PCR was performed in technical triplicate on a StepOnePlus™ Real-Time PCR System (Applied Biosystems, Cat# 4376600) using the Power SYBR® Green PCR Master Mix (Applied Biosystems, 4367660). Target gene expression was normalized to the average of two housekeeping genes (Gapdh and Rn18s), and fold change was calculated utilizing the ΔΔ-Ct method. Primers are listed in Table below.

#### Nucleotide sequence of primers used in this

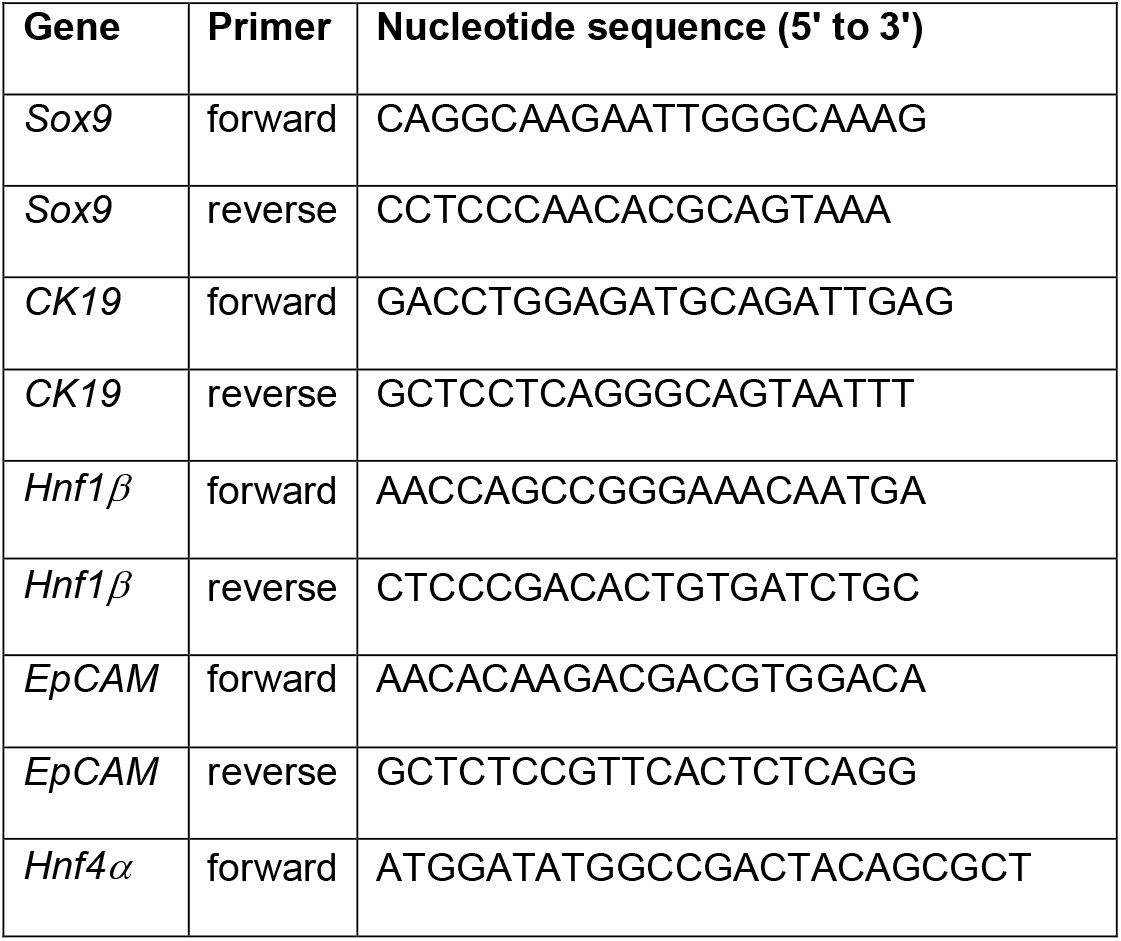
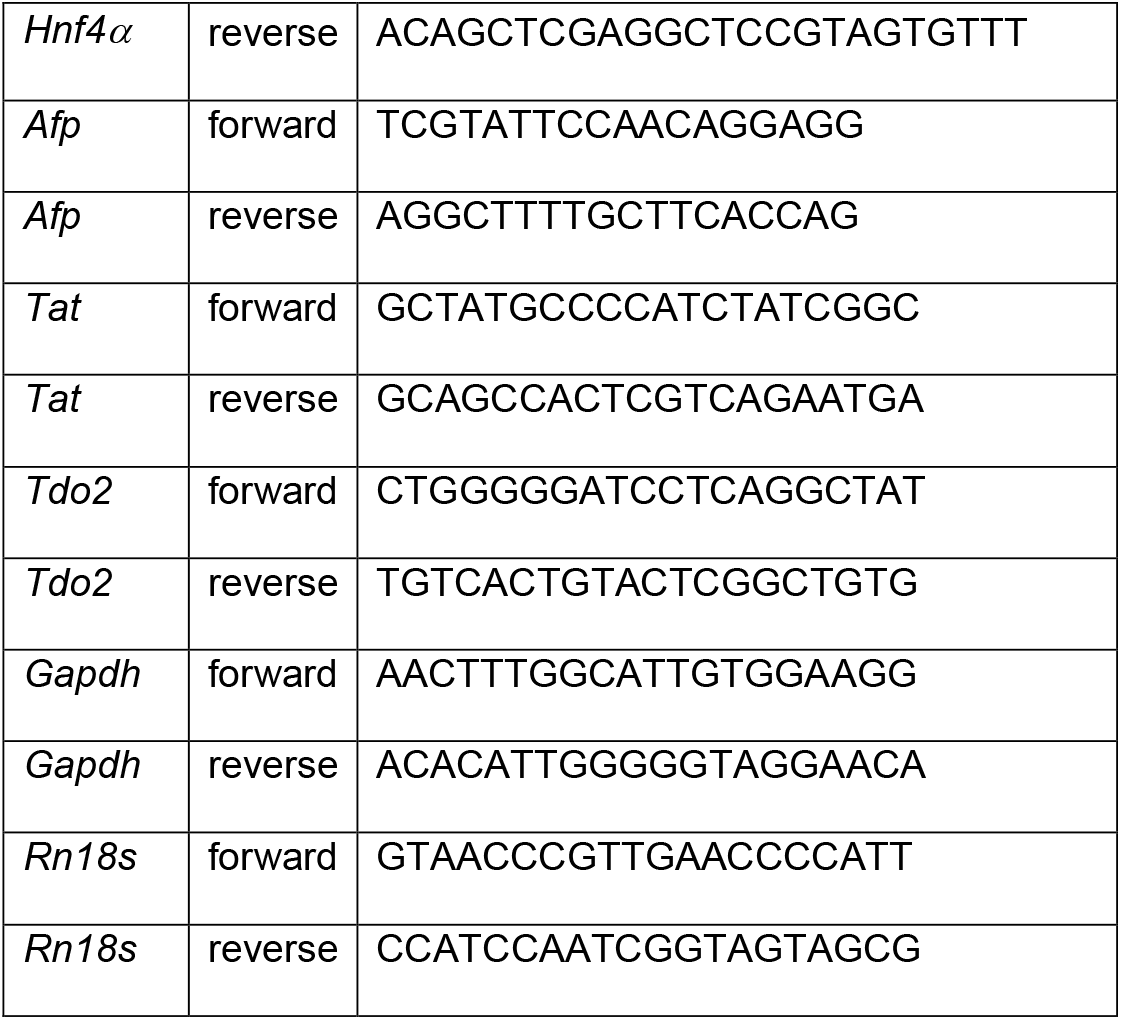

#### RNA-Seq Analysis

For each group (*Akt-YAP1 Sox9* WT and *Akt-YAP1 Sox9* LKO), two mouse liver samples were processed for RNA-seq analysis. RNA samples from livers were used to generate library using TruSeq kit from Illumina ^34, 35^. We used in-house HiSeq2500 platform and sequenced 200 million reads to accurately quantify genes & transcripts ^36^. Raw sequencing data was analyzed by FastQC for quality control ^37^. Low quality reads or adapter sequences were trimmed out by Trimmomatic ^38^. After pre-processing, sequenced reads were aligned to mouse reference genome mm10 by HISAT2 aligner ^39^. Read counts for each gene were then quantified by HTSeq ^40^. All the pipelines were run by default parameter settings. RNA-seq data have been submitted to the online database gene expression omnibus (GEO) accession ID: GSE200472. Differential expression (DE) analysis was performed to compare *Akt-YAP1 Sox9* WT versus *Akt-YAP1 Sox9* LKO. Based on the read counts, DE tests were performed by R package ‘DEseq2’ ^41^ and top DE genes were selected by absolute fold-change greater than 1 and FDR=0.05. These DE genes were then used as input for Ingenuity Pathway Analysis (IPA)® to call pathways that were significantly enriched with FDR=0.1.

#### Bioinformatic comparison between mouse and human models

To further investigate how the mouse models may mimic a subset of human liver cancer including CCA or HCC, public human transcript data were collected to compare with the gene signature obtained from the mouse models. For HCC, LIHC TCGA database was assessed similarly. After pre-processing, gene expression data were analyzed by R package ‘limma’ test ^42^ and top differentially expressed genes were selected by absolute fold-change greater than 1 and FDR=0.05 (same criteria as the mouse model). Top DE genes from human study were further used to detect significantly enriched pathways by IPA® software with FDR=0.1. To test the molecular similarity between our mouse *Akt-YAP1* liver cancer model and the human studies, three publicly available human CCA the cancer genome atlas program (TCGA) dataset was analyzed by three comparisons. (1) Pathway enrichment analysis. Top enriched pathways detected by mouse model and human studies were detected independently and compared. (2) Gene signature analysis. Top DE genes from mouse models were converted to human homologous genes by Mouse Genome Database (MGD) ^43^. These genes were then applied into the human studies to check their expression signatures. (3) Signature prediction. Positive prediction of gene signatures tested in this study has been calculated by Nearest Template Prediction (Gene Pattern module), ^44^ as previously described ^15^. For CCA studies, each patient was also assigned to either the Proliferation or Inflammation class based on the results of unsupervised clustering as previously described ^45^. Additional signatures assessed were for Notch activation ^46^ and hepatic stem-cell like group of ICC patients ^47^. Any significant correlation to a subclass was noted by p value (p<0.05).

### Statistical Analysis

For all mouse experiments, sample size was pre-determined based on previous literature describing SB-HDTVI-mediated liver carcinogenesis ^32^. Accordingly, littermates were randomized into groups for HDTVI and managed throughout the course of treatment in a non-blinded manner. All subsequent molecular, immunohistochemical, and immunofluorescence analysis was performed in a blinded manner. All confidence intervals shown on the bar plots are presented as mean ± standard error of mean (SEM). Differences in mean values of liver volume and LW/BW ratio were analyzed by one-way ANOVA assuming normal Gaussian distribution with Geisser Greenhouse posttest correction. p<0.05 was considered significant (*), p<0.01 was considered highly significant (**), p<0.005 was considered extremely significant (***), and so on. All statistical analysis on patient samples has been included in the results section and respective p-values were included in the pertinent text and figure legends. All statistics were performed using GraphPad Prism 10.0 (GraphPad Software) or R software.

## Acknowledgement

We extend special thanks to Dr. Daniela Sia at Icahn School of Medicine for her kind guidance in bioinformatic analysis.

## Notes

**Funding:** This work was supported by NIH grants R01CA258449 NIH grant to S.K. and 1P30DK120531-01 to Pittsburgh Liver Research Center (PLRC) and Innovation in Cancer Informatics Discover grant (https://www.the-ici-fund.org) to S.K and S.L., and partly by R01 DK130949 to M.O.

**Conflict of Interest:** There are no financial conflicts of interest to declare relevant to the current manuscript for any of the authors.

### Competing Interest Statement

The authors have declared no competing interest.

